# Mitotic spindle positioning protein (MISP) is an actin bundler that senses ADP-actin and binds near the pointed ends of filaments

**DOI:** 10.1101/2023.05.05.539649

**Authors:** E. Angelo Morales, Matthew J. Tyska

## Abstract

Actin bundling proteins crosslink filaments into polarized structures that shape and support membrane protrusions including filopodia, microvilli, and stereocilia. In the case of epithelial microvilli, mitotic spindle positioning protein (MISP) is an actin bundler that localizes specifically to the basal rootlets, where the pointed ends of core bundle filaments converge. Previous studies established that MISP is prevented from binding more distal segments of the core bundle by competition with other actin binding proteins. Yet whether MISP holds a preference for binding directly to rootlet actin remains an open question. Using *in vitro* TIRF microscopy assays, we found that MISP exhibits a clear binding preference for filaments enriched in ADP-actin monomers. Consistent with this, assays with actively growing actin filaments revealed that MISP binds at or near their pointed ends. Moreover, although substrate attached MISP assembles filament bundles in parallel and antiparallel configurations, in solution MISP assembles parallel bundles consisting of multiple filaments exhibiting uniform polarity. These discoveries highlight nucleotide state sensing as a mechanism for sorting actin bundlers along filaments and driving their accumulation near filament ends. Such localized binding might drive parallel bundle formation and/or locally modulate bundle mechanical properties in microvilli and related protrusions.

## INTRODUCTION

Actin filament assemblies provide structural support for a variety of cell surface features, which mediate interactions with the external environment in diverse biological contexts. Two general architectures include actin meshworks and bundles; meshworks generate large mechanical forces, such as those needed for leading edge protrusion during cell motility, whereas bundles generate localized forces that can power the extension of fingerlike membrane protrusions.^1,2^ Examples of the latter case include filopodia, which extend from crawling cells to promote surface attachment and control steering.^3^ Other well-studied protrusions that are supported by actin bundles include microvilli, which extend from the apex of solute-transporting epithelial cells,^4,5^ and morphologically related stereocilia, which comprise the mechanosensory hair bundle that resides at the functional surface of cochlear and vestibular hair cells.^6^ In all three of these cases, individual protrusions are supported by a core bundle consisting of multiple actin filaments, ranging from ~25 for microvilli up to 100s for stereocilia.^7–9^ Multiple filaments are needed because individual polymers exhibit a flexural rigidity that is too low to support plasma membrane deformation.^10^ By bundling multiple filaments together with cell-type specific factors, cells create more rigid structures that promote and maintain membrane protrusion without buckling.

A defining feature of a protrusion core bundle is the uniform polarity of its constituent actin filaments. In all cases, actin filament barbed ends are oriented out into the distal tip of the protrusion, whereas the pointed ends extend down into the cytoplasm.^11,12^ Importantly, the critical concentration for incorporation of ATP-actin (the most abundant form of monomer in the cytoplasm) is much lower at the barbed relative to the pointed end (0.12 vs. 0.6 µM, respectively).^13^ The differential kinetic properties at the two ends fuel dynamic behaviors such as treadmilling, where new ATP-actin monomers incorporate at the barbed end, flux through the polymer where hydrolysis occurs, and then exit from the pointed ends in the ADP-bound form.^14^ Filament networks that exhibit treadmilling also generate mechanical force at their growing barbed ends, which can be harnessed to power subcellular and cell scale activities ranging from the motion of vesicles and endomembranes to the leading-edge protrusion that powers cell motility.^15,16^ Treadmilling has also been observed in core bundles that support fingerlike protrusions. For instance, filopodia are supported by core bundles that exhibit robust treadmilling,^17^ and newly formed microvilli on differentiating epithelial cells also demonstrate treadmilling,^18,19^ which powers their gliding motility across the apical surface.^20^ How the behaviors of individual filaments in treadmilling parallel actin bundles are coordinated remains unclear, although *in vitro* studies implicate factors with multivalent filament binding potential, such as VASP at filament barbed ends.^21^

In addition to creating a kinetic scenario that supports treadmilling, the eventual hydrolysis of ATP in newly incorporated monomers at the barbed end creates a gradient of nucleotide states consisting of ATP-, ADP-Pi-, and ADP-bound actin along the length of an actively growing filament.^22^ This gradient in nucleotide composition also impacts the physical properties of the filament, such as flexibility, which is increased in older polymer that is rich in ADP-actin.^23,24^ The nucleotide landscape of an actin filament might also impact where regulatory and network building proteins are able to bind. One established example of this type of “sorting” is found in studies of cofilin, a ubiquitously expressed filament severing protein that only binds to older ADP-actin.^25^ In contrast, coronin-1B displays a binding preference for ATP and ADP-Pi-bound actin.^26,27^ Another well-studied factor is the Arp2/3 complex, which crosslinks newly polymerizing filaments to the sides of pre-existing filaments in the ATP and ADP-Pi states.^28^ Although the nucleotide-dependent targeting and function of these factors have provided a mechanistic framework to understand how branched actin networks are built and recycled,^1^ whether actin nucleotide states also control the formation and organization of parallel bundles, like those that shape membrane protrusions, remain unexplored.

Interestingly, the bundling proteins that comprise microvillar core bundles - fimbrin, espin, villin, and MISP – do localize to different segments along the length of the structure.^29,30^ While espin and villin localize along the full length of the core bundle, as expected for canonical bundlers,^31,32^ fimbrin seems to prefer the basal half.^29^ Unlike these bundlers, MISP exclusively decorates the rootlets.^30^ Although the selective localization of MISP is partially dependent on ezrin’s membrane-actin crosslinking activity,^30^ whether intrinsic features of core bundle actin filaments, such as monomer nucleotide state, contribute to the sorting of MISP or other bundlers remains an open question.

In this study, we report that MISP exhibits an intrinsic ability for binding directly to rootlet actin. Using *in vitro* reconstitution assays coupled with total internal reflection fluorescence (TIRF) microscopy, we found that MISP demonstrates a clear binding preference for filaments enriched in ADP-actin, and assays with actively growing actin filaments revealed that MISP binds at or near their pointed ends. Bundling assays also revealed that MISP can organize multiple actin filaments into a parallel bundle, a feature that is likely conferred by its pointed end binding ability. These findings implicate sensing of actin nucleotide states in the specific targeting of MISP to microvillar rootlets, and further suggest that MISP may play a role in organizing nascent filaments in a parallel fashion at early stages of microvillar assembly.

## RESULTS

### Microvillar core bundle rootlets are enriched in ADP-actin

In cultured epithelial model systems, the core actin bundles that support nascent microvilli undergo treadmilling,^18–20^ and thus their rootlets are expected to be enriched in ADP-actin. To determine if the rootlets of mature microvilli in native tissue are also enriched in ADP-actin, we immunostained mouse small intestinal tissue with antibodies targeting cofilin, a severing protein that specifically binds to actin in the ADP-bound state.^25,33^ Confocal microscopy of the resulting tissue sections revealed that cofilin is highly enriched on microvillar rootlets in a pattern that closely resembles MISP localization as we previously reported (Figure S1A,B).^30^ Thus, *in vivo* MISP binds to filaments that appear to be enriched in ADP-actin.

### MISP dissociates slowly from ADP-actin enriched filaments

Strong colocalization of cofilin and MISP on microvillar rootlets suggests that MISP may prefer binding to filaments enriched in ADP-actin monomers. To directly test this possibility, we turned to *in vitro* reconstitution assays with purified proteins using TIRF microscopy. We purified a MBP-EGFP tagged version of MISP (herein referred to as ‘MISP’) as previously described^30^ and generated biotin-rhodamine-labeled actin filaments enriched with monomers in three distinct nucleotide states. ADP-actin and ADP-Pi-actin filaments were generated using phalloidin as previously described (Figure 1A,B; top row).^34^ To mimic the ATP state, we used AMP-PNP as a nucleotide analog (Figure 1C; top row). Importantly, we previously showed that phalloidin labeling does not disrupt MISP’s interaction with filaments.^30^ To determine if MISP’s dissociation from filaments is impacted by monomer nucleotide state, MISP was introduced at low concentrations (16.5 nM) to surface bound filaments in each nucleotide state and binding event lifetimes were measured directly using kymography. Notably, MISP dwell times on ADP-actin enriched filaments were significantly longer relative to filaments enriched in ADP-Pi- or AMP-PNP-actin, respectively (10.72 s [95% CI = 10.15 – 11.33]; 6.78 s [95% CI = 6.13 – 7.50]; 5.83 s [95% CI = 5.53 – 6.14]) (Figure 1A-F; S1F). Consistently, the number of events longer than 60 seconds was also significantly higher in the ADP condition relative to ADP-Pi and AMP-PNP (Figure S1G). We note here that the measured dwell times are likely underestimates as we did not consider events lasting longer than our imaging timeframe of 500 sec. Nevertheless, these data indicate that MISP dissociation from filaments containing ADP-actin is slow relative to other nucleotide states. To determine if MISP’s association to filaments is also dependent on nucleotide composition, we monitored the rate of increase in MISP fluorescence intensity during the decoration of immobilized actin filaments. Exponential fitting of the resulting time courses revealed that the apparent association rate is slightly lower for ADP- vs. ADP-Pi- and AMP-PNP-actin containing filaments, by factors of ~1.1 and ~1.2, respectively (0.30 µM^-1^s^-1^ vs. 0.34 µM^-1^s^-1^ vs. 0.36 µM^-1^s^-1^) (Figures S1C-E). Thus, MISP’s association kinetics are unlikely to contribute to its preferential occupancy on ADP-actin enriched filaments. Together, these *in vitro* findings suggest that an actin nucleotide state-sensing mechanism may contribute to MISP’s specific enrichment on microvillar rootlets.

**Figure 1.**
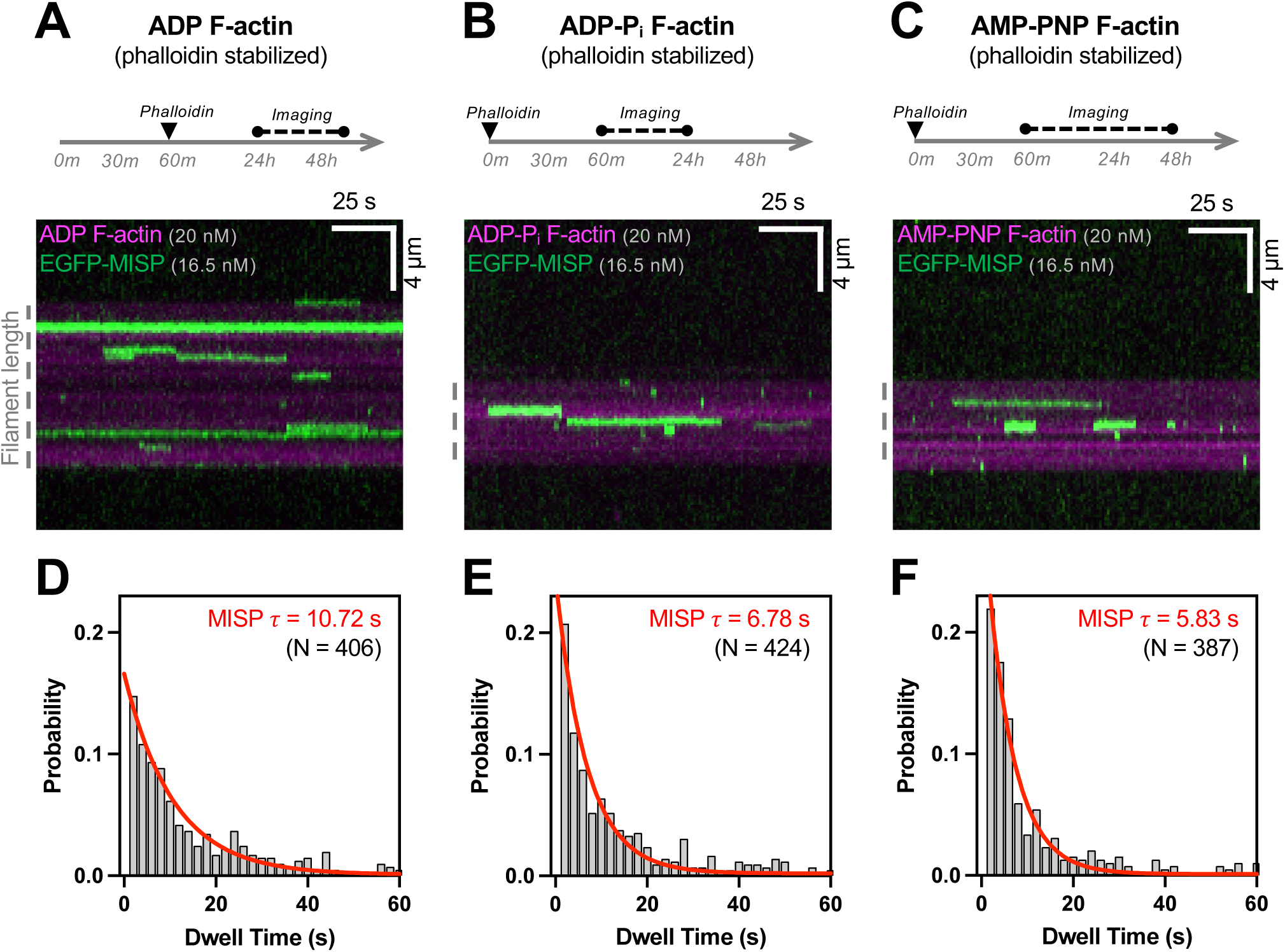
MISP preferentially binds to ADP-bound actin filaments. **(A-C)** Top row: Experimental setup showing the preparation and imaging of F-actin in each nucleotide state: ADP (A), ADP-Pi (B), AMP-PNP (C). In all conditions, actin polymerization begins at t = 0; Bottom row: TIRF microscopy kymographs representative of dwell times of EGFP-MISP (green) on each indicated nucleotide state of rhodamine F-actin (magenta). Dashed gray line next to each panel denotes the length of filament. **(D-F)** Probability distribution of all MISP dwell times in ADP F-actin (D), ADP-Pi F-actin (E) and AMP-PNP F-actin (F). Bin size = 2 s. Non-linear one-phase exponential decay fittings for each distribution are shown in red. The X axis (i.e., dwell time) was purposefully ended at 60 s to display comparisons between fitting curves at early time points. All data in each condition are representative of at least three independent experiments.

### MISP preferentially binds near the pointed ends of actively growing actin filaments

We next sought to determine if MISP exhibits preferential binding near the pointed ends of actively growing filaments, as we expect these sites to be enriched in ADP-actin after several minutes of polymerization.^35^ To this end, we designed TIRF assays with actively growing filaments and observed polymerization for ~6 min prior to the addition of MISP, which enabled identification of barbed and pointed ends based on differences in their growth rates (Figure 2A). After flowing in low concentrations of MISP, we observed puncta preferentially binding with long dwell times (> 70 s) at or near the pointed ends (Figure 2B-E; Video S1). In contrast, we observed fewer binding events near elongating barbed ends, which are more likely to be enriched in ATP- or ADP-Pi-actin (Figure 2E). Interestingly, dwell times for MISP end-binding events were much longer than side-binding events (Figure 2F). To determine if this difference in event lifetime is based on nucleotide state differences between these sites, we conducted experiments like those described above (Figures 1 and S1) by introducing low concentrations of MISP to phalloidin-stabilized filaments containing only ADP-actin (Figure S2A). Even under these conditions, however, the number of MISP end-binding events was significantly higher compared to side-binding events across multiple fields of view (Figure S2B-C).

**Figure 2.**
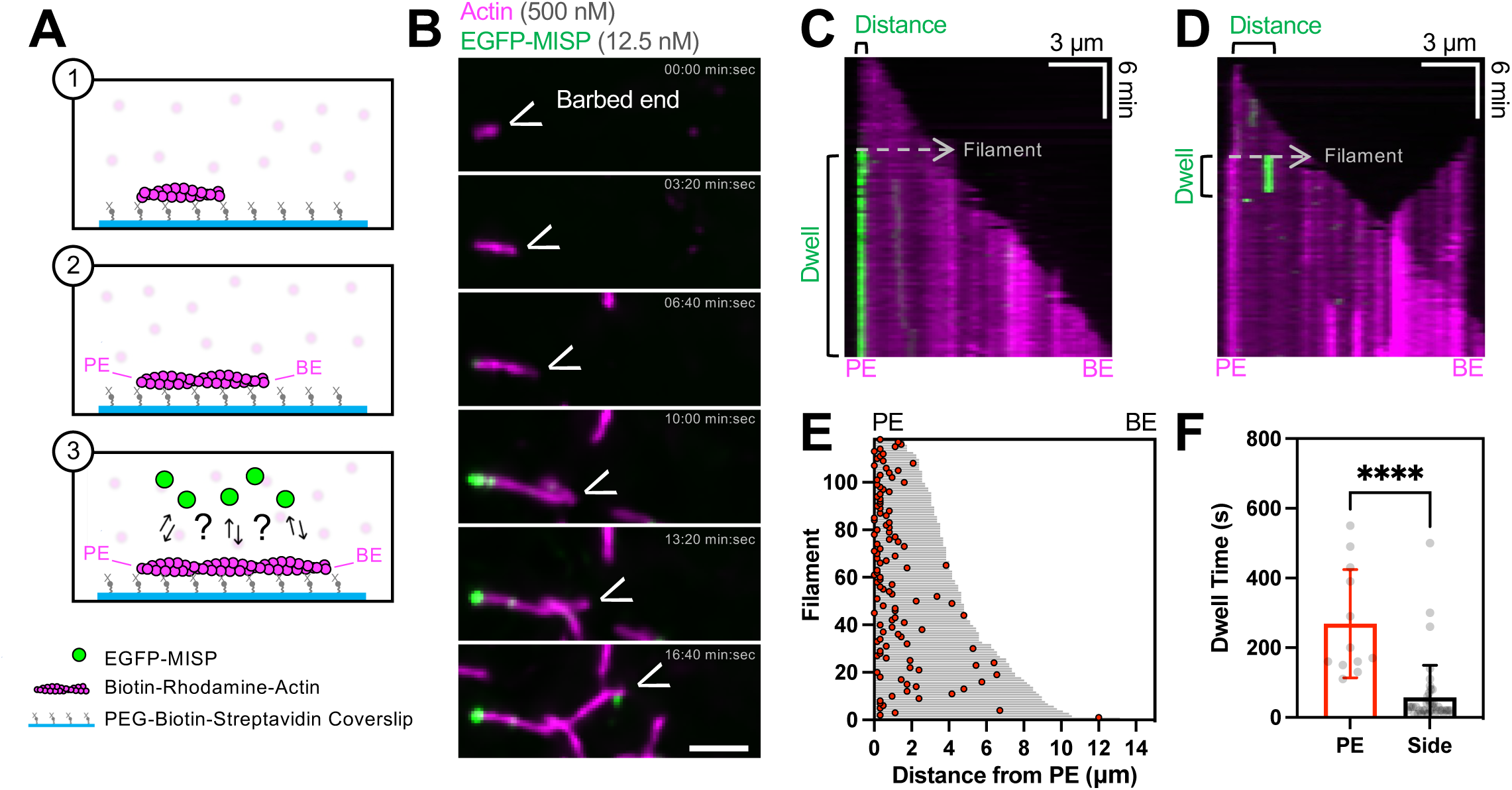
MISP preferentially binds near the pointed ends of actin filaments. **(A)** Cartoon schematic of the experimental setup (1-2) and possible outcomes (3). **(B)** TIRF microscopy montage of a tethered polymerizing rhodamine F-actin (magenta) and EGFP-MISP molecules (green). Scale bar = 3 µm. **(C-D)** TIRF microscopy kymographs representative of EGFP-MISP’s binding events (green) on F-actin (magenta), near their pointed end (C), and at their side (D). **(E)** Distribution of EGFP-MISP’s binding events on actin filaments of known polarity. Each filament length is plotted as a horizontal bar in gray, and the distance of each binding event from the pointed ends (“PE”) is plotted as a red dot. **(F)** Mean dwell time of MISP’s binding events near the pointed ends (“PE”) or at the filament side (“Side”). Each dot represents a single event. Bar plots and error bars denote mean ± SD. p value was calculated using the unpaired t test (****p < 0.0001). All data are representative of three independent experiments.

In cultured intestinal epithelial cells, MISP appears to be anchored to pre-existing actin networks before core bundles assemble.^30^ From this perspective, we sought to determine if substrate bound MISP can capture the pointed ends of newly polymerizing filaments from solution. To examine this possibility, we tethered MISP molecules on the coverslip using anti-MISP antibodies and flowed in short seeds of actively polymerizing actin filaments (Figure 3A). Remarkably, immobilized MISP molecules maintained preferential binding near the pointed ends, as indicated by the fast elongation of the opposite barbed end over several minutes (Figure 3B-C; Video S2). When binding events were binned as pointed end, barbed end, or side-binding, we again found a preference for pointed end binding (Figure 3D). Together these data indicate that MISP exhibits a strong preference for ADP-actin and for direct end binding, which likely work in combination to constrain this factor at or near filament pointed ends.

**Figure 3.**
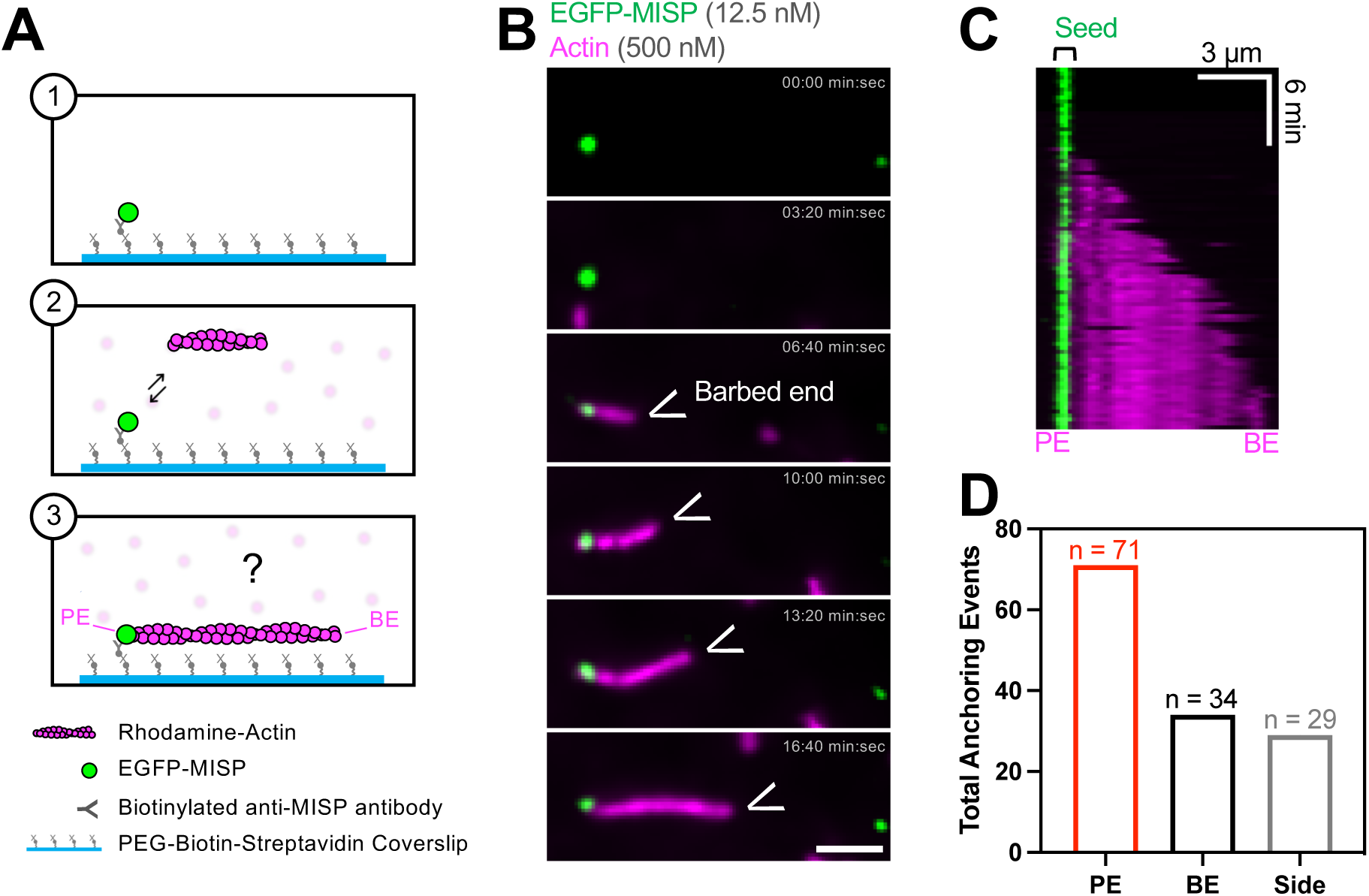
MISP preferentially captures actin filaments from the pointed ends. **(A)** Cartoon schematic of the experimental setup (1-2) and predicted outcome (3). **(B)** TIRF microscopy montage of immobilized EGFP-MISP molecules (green) anchoring a polymerizing rhodamine F-actin (magenta) from the pointed end. Scale bar = 3 µm. **(C)** Kymograph of movie corresponding to the montage in (B). **(D)** Quantification of EGFP-MISP-driven anchoring events from the pointed ends (“PE”), barbed ends (“BE”), and side (“Side”) of actin filaments. All data are representative of three independent experiments.

### MISP drives parallel and anti-parallel bundling of surface bound filaments

Considering that MISP normally functions as a crosslinker of multiple filaments,^30,36^ we next investigated whether MISP’s binding preference for ADP-actin and filament pointed ends impacts its ability to organize multi-filament bundles. One intriguing possibility is that MISP uses its ADP-actin binding ability to sequester the pointed ends of filaments, to in turn create bundles of uniform polarity. To test this idea *in vitro*, we combined untethered, polymerizing actin filaments with soluble MISP, and then observed the dynamics of bundle formation over time using TIRF microscopy. Visualizing two-filament bundling events, we found that MISP is capable of bundling filaments in parallel and antiparallel fashion, with no significant preference for a single orientation (Figure 4A-C; Video S3). We also noted that MISP intensity was higher in two-filament bundles compared to single filaments (Figure 4B, D), suggesting a stronger affinity for two-filament arrays, likely reflecting MISP’s multivalent actin binding potential.^36^ A close inspection of two-filament parallel bundling events also revealed that MISP signal was lower on newly polymerizing pairs of filaments (Figure 4B; top row, green channel), as expected based on its preference for binding older filaments that are enriched in ADP-actin (Figure 1C).

**Figure 4.**
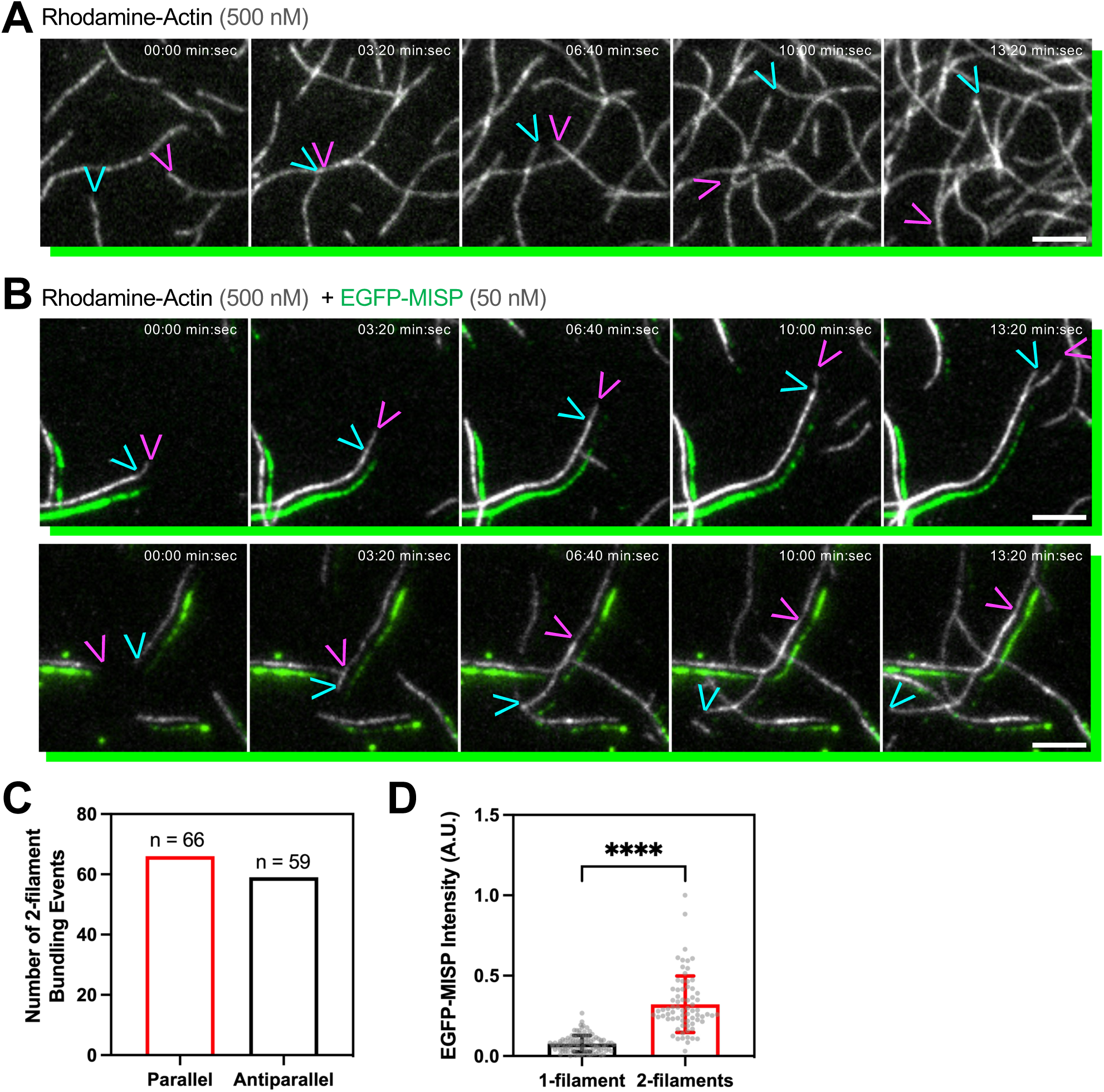
MISP bundles actin filaments in a parallel and antiparallel manner. **(A-B)** TIRF microscopy montages of rhodamine F-actin (gray), and EGFP-MISP (green). (A) Control experiment without EGFP-MISP. (B) 2-filament bundling event driven by EGFP-MISP in a parallel (top row) and antiparallel (bottom row) manner. Green channels in A and B were slightly shifted to the lower right side of image to better visualize the overlap between F-actin and MISP. Colored arrowheads indicate the position of barbed ends in each time point. Scale bar = 5 µm. **(C)** Quantification of 2-filament parallel and antiparallel bundling events from movies as shown in (B). All data are representative of at least three independent experiments. **(D)** Mean intensity of EGFP-MISP on 1-filament versus 2-filaments from movies as shown in (B). Bar plots and error bars denote mean ± SD. p value was calculated using the unpaired t test (****p < 0.0001).

### MISP drives parallel alignment of multi-filament bundles in solution

During the course of collecting timelapse datasets in the experiments outlined above, we also observed MISP-decorated actin bundles of four or more filaments (based on intensity) landing in the field of view (Figure S3A-C; Video S4, center panel). If MISP exerts no influence over the orientation of individual polymers in multi-filament bundles, one might expect to only observe anti-parallel bundles as the number of filaments in a bundle increases (Figure S3B; top row). However, we observed thicker bundles that landed and then elongated their barbed ends uniformly in one direction, implying parallel alignment of constituent filaments (Figure S3B; bottom row). A close inspection across multiple imaging fields revealed broom-shaped actin structures consisting of 20 or more filaments and strong MISP signal at one end, which is consistent with parallel bundling (Figure S3C). Our failure to observe parallel bundling events *de novo* on the surface of the coverslips (Figure 4) is most likely due to the reduced conformational flexibility imposed by substrate attachment. Together, these results suggest that MISP holds the capacity to create anti-parallel and parallel bundles of multiple filaments. However, the increased conformational degrees of freedom offered in solution may allow MISP to sequester ADP-actin enriched pointed ends and promote parallel bundle formation.

## DISCUSSION

MISP localization at microvillar rootlets is required for the normal elongation of this end of the core bundle down into the sub-apical terminal web.^30^ Confinement of MISP to core bundle rootlets is partially explained by its exclusion from more distal segments by the membrane-cytoskeleton crosslinking activity of ezrin.^30^ The unique MISP actin binding properties identified in this work represent additional, complementary mechanisms for targeting MISP specifically to rootlets. Using *in vitro* reconstitution assays and direct visualization with TIRF microscopy, we found that MISP dwell times on actin filaments are significantly increased by the presence of ADP-actin relative to other nucleotide states (ADP-Pi and ATP-like AMP-PNP) (Figure 1). MISP also demonstrated a clear preference for binding near the pointed ends, even in filaments that were composed entirely of ADP-actin. Here, we use the word “near” as the diffraction limited nature of TIRF microscopy does not allow us to discriminate between true capping vs. binding in close proximity to the pointed ends. Nevertheless, because core bundle rootlets are enriched in ADP-actin (Figure S1) and the pointed ends of constituent actin filaments,^11^ these *in vitro* results provide a mechanistic explanation for the direct targeting of MISP to core bundle rootlets.

Recent ultrastructural studies showed that ADP-actin containing filaments display a flexibility that may expose multiple binding sites beyond those unmasked by phosphate release alone,^24,37^ and this may further permit the association of factors that appear to “sense” nucleotide states, such as MISP. In terms of the molecular nature of the MISP-actin interactions, the mechanism is likely unique as MISP shares little homology with cofilin or other factors that demonstrate a preference for binding at or near the pointed ends of actin filaments, including tropomodulins,^38^ CAP, emerin,^39^ and the bacterial effector VopL/F.^40^ Additional studies on the mechanism that allows MISP to sense ADP-actin or bind near the pointed ends, will be needed to fill in these important mechanistic details.

What functional benefit(s) would ADP-actin sensing and pointed end binding provide for a filament bundling protein such as MISP? Although it currently remains unclear how MISP drives rootlet elongation, interestingly possibilities emerge considering the ADP-actin sensing and pointed end binding activities revealed here. For example, MISP might promote monomer association at the pointed ends, potentially by competing off capping proteins such as tropomodulin-3.^41,42^ The general idea that a bundling protein might regulate the activity of an end binder is supported by recent *in vitro* biochemical studies showing that espin exerts anti-capping effects at the barbed ends, which ultimately contribute to filament and bundle elongation.^43^ Alternatively, MISP could prevent monomer dissociation from the pointed ends, potentially by delaying cofilin-driven severing. The specific localization of MISP to core bundle rootlets might also drive a downstream sorting mechanism that favors (or restricts) the binding of other actin regulatory factors along the axis of the core bundle.^44^ Previous studies found that different bundling proteins self-segregate within the same actin networks, which may further promote the sorting of other actin-binding factors by competition or by the differential filament interspacing they create.^44,45^ Sorting was also demonstrated in previous studies with fimbrin, which displays preferential binding to MISP-bundled filaments, as well as the basal half of the core bundle.^29,30^ Although these roles are speculative, MISP is well positioned to participate in these activities, alone or in combination, to promote rootlet elongation.

Beyond regulating the length of core bundle rootlets at steady-state (as alluded to above), ADP-actin sensing and pointed end binding might also help MISP coordinate the organization of actin filaments during core bundle assembly and microvillar growth. In the case of filopodia, supporting core bundle filaments have their pointed ends crosslinked to the sides of pre-existing filaments via the Arp2/3 complex, and this meshwork serves as an anchoring platform for orienting filaments during protrusion growth.^46^ Given that microvilli assemble in an Arp2/3-independent fashion,^47,48^ it remains unclear how the uniform polarity of core bundle filaments is established. Our bundling assays indicate that MISP assembles substrate associated F-actin in both parallel and antiparallel bundles (Figure 4), which is consistent with MISP localization to the unipolar core bundles of microvilli and the mixed polarity bundles of stress fibers in cultured cells.^30,36^ However, the broom-shaped structures that we observed falling out of solution and landing on the TIRF surface are suggestive of multi-filament bundles with uniform polarity (Figure S3). Fimbrin assembles parallel bundles *in vitro*,^49^ but its recruitment to microvillar rootlets seems to be MISP-dependent.^30^ Espin is another microvillar bundler capable of creating bundles with uniform polarity in cells,^50^ although its late accumulation during microvilli differentiation is at odds with orchestrating bundle polarity.^31^ Thus, as an early microvillar factor with the ability to preferentially bind and crosslink filaments near their pointed ends, MISP may be a candidate for establishing the polarity of core bundles early in microvillar assembly.

Although the data we present here suggests that MISP might promote the uniform polarity of filaments during bundle formation *in vitro*, in nascent epithelial cells this factor likely coordinates with other actin binding and bundling factors that are known to be present during the earliest stages of microvillar growth. One such factor is Epidermal Growth Factor Receptor Pathway Substrate 8 (EPS8), a distal tip-specific microvillar protein that also holds bundling potential *in vitro*.^51–53^ Like MISP, EPS8 strongly localizes to the apical region of immature enterocytes within intestinal crypts.^30,52^ Given their end-specific targeting and bundling functions, it is tempting to speculate that EPS8 and MISP orchestrate the polarity of nascent core bundles from the barbed and pointed ends, respectively. Assays that allow direct visualization of microvillar growth will be required to determine whether these factors colocalize at the right place and time to organize polymerizing filaments into parallel core actin bundles.^52^

## Supporting information

Morales and Tyska Supp Figs

Morales and Tyska VideoS2

Morales and Tyska VideoS1

Morales and Tyska VideoS3

Morales and Tyska VideoS4

## ACKNOWLEDGMENTS

The authors would like to thank all members of the Tyska laboratory for their constructive feedback. Microscopy was performed in part through the VUMC Cell Imaging Shared Resource. This work was supported by the NIH grants DK095811, DK125546, DK111949 (M.J.T.).

## AUTHOR CONTRIBUTIONS

Conceptualization, E.A.M. and M.J.T.; Methodology, E.A.M. and M.J.T.; Validation, E.A.M.; Formal Analysis, E.A.M.; Investigation, E.A.M.; Writing, E.A.M. and M.J.T.; Visualization, E.A.M.; Supervision, M.J.T.; Project Administration, M.J.T.; Funding Acquisition, M.J.T..

## INCLUSION AND DIVERSITY

We support inclusive, diverse, and equitable conduct of research.

## SUPPLEMENTAL INFORMATION

**Figure S1. MISP preferentially binds to aged actin filaments *in vivo* and *in vitro*. Related to Figure 1. (A-B)** Confocal images of separate frozen small intestinal sections stained for membrane with WGA (blue), F-actin with phalloidin (magenta), cofilin (yellow; panel A), and MISP (yellow; panel B). Each panel shows a split two-color merge. Bottom rows show inverted single channels for each marker from in A and B. Scale bar = 10 µm. **(C-E)** Fluorescence intensity of EGFP-MISP (50 nM) on immobilized F-actin (50 nM) in each indicated nucleotide state as a function of time. Non-linear exponential fittings shown in red were used to determine the apparent association rates for each condition. All data in each condition are representative of three independent experiments. k_off_: dissociation constant; k_on-app_: apparent association constant **(F)** Time constants of EGFP-MISP for each replicate used in Figure 1D-F. **(G)** Number of EGFP-MISP binding events lasting longer than 60 seconds in each nucleotide state of actin.

**Figure S2. MISP preferentially binds to the ends of stabilized ADP-actin filaments. Related to Figure 2. (A)** Cartoon schematic of the experimental setup (1-2). **(B)** TIRF microscopy image of phalloidin-stabilized biotin-rhodamine-actin filaments (magenta), and EGFP-MISP molecules (green). Scale bar = 5 µm. **(C)** Single binding events of MISP at the ends (“End”) or at the side (“Side”) of stabilized actin filaments from B. Each dot represents a single event. Bar plots and error bars denote mean ± SD. p value was calculated using the unpaired t test (***p < 0.001).

**Figure S3. MISP assembles multi-filament parallel and antiparallel bundles. Related to Figure 4. (A-C)** TIRF microscopy montages of biotin-rhodamine F-actin (black), and EGFP-MISP (green). (A) Control experiment without EGFP-MISP. (B) Multi-filament antiparallel (middle row) and parallel bundling (bottom row) events driven by EGFP-MISP; AP = antiparallel bundling event; P = parallel bundling event. (C) Examples of multi-filament parallel bundling events before and after polymerization of their surface-bound actin filaments; P = parallel bundling event. Scale bar = 10 µm.

## MOVIE LEGENDS

**Video S1. MISP preferentially binds near the pointed ends of actin filaments. Related to Figure 2B.** TIRF microscopy movie of freely diffusing EGFP-MISP (green) preferentially binding near the pointed ends of a tethered polymerizing actin filament (magenta). Scale bar = 3 µm.

**Video S2. MISP preferentially captures actin filaments from the pointed ends. Related to Figure 3B.** TIRF microscopy movie of immobilized EGFP-MISP (green) anchoring a freely diffusing actin filament near its pointed end. Scale bar = 3 µm.

**Video S3. MISP forms two filament bundles in parallel and antiparallel fashion. Related to Figure 4.** TIRF microscopy movie of MISP-driven bundling of two actin filaments (white). Left panel shows controls with no MISP. Middle and right panels show conditions with MISP, resulting in parallel and antiparallel bundling events, respectively. Scale bar = 4 µm.

**Video S4. MISP bundles multiple actin filaments in parallel and antiparallel configurations. Related to Figure S4.** TIRF microscopy movie of MISP-driven bundling of multiple actin filaments (black) landing on the field of view. Left panel shows control with no MISP. Middle and right panels show conditions with MISP, which result in parallel and antiparallel bundling events, respectively. Scale bar = 4 µm.

## MATERIAL AND METHODS

### Tissue immunostaining

Frozen tissue sections were washed with 1X PBS and blocked with 5% Normal Goat Serum (Abcam; ab7481) for 2h at room temperature. Sections were washed and incubated with primary antibodies overnight at 4°C. Primary antibodies used were rabbit anti-MISP (Thermo Scientific; PA5-61995), or rabbit anti-Cofilin (Sigma; C8736). Tissue sections were then washed with 1X PBS and incubated with Alexa-Fluor-568 Phalloidin (Invitrogen; A12380), Wheat German Agglutinin (WGA) 405M (Biotium; 29028-1), and F(ab’)2-goat ant-rabbit IgG Alexa-Fluor-488 (Molecular probes; A11070) for 2 h at room temperature. Tissue sections were washed with 1X PBS and mounted in ProLong Gold (Invitrogen; P36930).

### Light microscopy and image processing

Total Internal Reflection Fluorescence (TIRF) and Laser Scanning Confocal microscopy imaging were conducted using the Nikon A1 Microscope equipped with 405, 488, 561 nm LASERs, Apo TIRF 100x/1.45 NA objective, and an Evolve EMCCD camera for the TIRF modality (Photometrics Technology). Images were denoised using Nikon Elements software for images shown in Figures 2, 3, S2 and S4. During imaging acquisition, the gain was matched between samples for comparison.

### MISP cloning and protein purification

The full length human MISP sequence was originally obtained from Harvard PlasmID Database (HsCD00326629). MISP was then shuttled into Gateway-adapted plasmid pEGFP-C1. To create baculovirus expression vectors, EGFP-MISP sequence was subcloned into modified pFastBac-6xHis-MBP LIC expression vector (Addgene; plasmid #30116). In-frame sequence insertion was confirmed by sequencing. The 6xHis-MBP-EGFP-MISP construct was expressed and purified from Sf9 insect cells as previously described.^30^ Briefly, insect cell pellets were resuspended in lysis buffer (20 mM Tris HCl, 0.3 M KCl, 10 mM imidazole, 10% glycerol, 2 mM DTT, pH 7.5) supplemented with protease inhibitors (Roche, 5892953001). The resultant lysate was then centrifuged at 35,000 rpm in a Ti 50.2 rotor (Beckman) for 30 minutes at 4 °C. Samples were loaded into a HisTrap column according to the manufacturer protocol and eluted with a 50-500 mM linear imidazole gradient (pH 7.5). Protein purity was assessed by SDS-PAGE. Eluted protein was concentrated using a centrifugal filter (Millipore; UFC803024) in storage buffer (20 mM Tris HCl, 0.1 M KCl, 10 mM imidazole, 10% glycerol, 1 mM EGTA, 2 mM DTT, pH 7.5).

### *In vitro* reconstitutions assays

#### Coverslip functionalization

Coverslips (Thorlabs, CG15KH) were placed into a glass jar, bathed in acetone for 30 minutes, incubated in ethanol for 15 minutes, and washed in MiliQ water three times. Subsequently, the coverslips were sonicated in 2% Hellmanex or 2% Micro90 for 2 hours at room temperature in a water bath sonicator, followed by series of washes in MiliQ water. Coverslips were then transferred into a 0.1 M KOH bath, incubated for 30 minutes, washed in MiliQ water and dried with clean nitrogen gas. For functionalization, coverslips were submerged in a glass jar containing mPEG-Silane (Laysan Bio Inc, MPEG-SIL-5000) or Biotin-mPEG-Silane (Laysan Bio inc, Biotin-PEG-SIL-5K) solution, and protected from light for 18 hours. The next day, coverslips were washed with clean ethanol and MiliQ water. Finally, coverslips were dried once again with clean nitrogen gas and stored at 4 °C for up to two weeks.

#### TIRF flow channel preparation

Strips of double-sided tape were placed along the length of the functionalized side of a coverslip leaving a gap of approximately 2-3 mm between each strip. Flow channels were made by placing the coverslip/tape strips on a clean glass slide. The flow channels were subsequently sealed by gently pressing the areas with tape in between glass sides, and stored at 4°C. Before each experiment, the flow channels were washed with high salt (50 mM Tris-HCl pH 7.5, 600 mM KCl, 1% BSA) and low salt (50 mM Tris-HCl pH 7.5, 150 mM KCl, 1% BSA) buffers, blocked with 1X TBSA (50 mM Tris-HCl pH 7.5, 50 mM KCl, 1% BSA) for 5 minutes, and coated with 0.1 – 1 μg/ml streptavidin (SIGMA, 189730) for 5 minutes.

#### Actin preparation

Rhodamine-actin (Cytoskeleton, AR05), biotin-actin (Cytoskeleton, AB07), and black actin (Cytoskeleton, AKL99) were resuspended in G-actin buffer (5 mM Tris-HCl, pH 8.0; 0.2 mM CaCl_2_) supplemented with 0.2 mM ATP (SIGMA, 10519979001) and mixed at a final ratio of 20:1:79, respectively. Alternatively, G-buffer was supplemented with 0.2 mM AMP-PNP (SIGMA, A2647) to prepare non-hydrolysable ATP-bound filaments. Subsequently, the resulting G-actin mix was ultracentrifuged at 90K rpm for 30 minutes at 4°C in an ultracentrifuge (Beckman, Optima TL 100) to remove actin oligomers and/or aggregates.

#### MISP binding on immobilized F-actin

G-actin was mixed with TIRF polymerization buffer (50 mM KCl, 1 mM MgCl_2_, 1 mM EGTA, 10 mM Imidazole pH 7.0, 50 mM DTT, 0.2 mM ATP, 15 mM Glucose, 0.5% Methylcellulose, 1X Oxyrase, 0.1% BSA), and flowed in through a streptavidin-coated flow channel at a final concentration of 500 nM. At this concentration, actin is expected to polymerize exclusively from the barbed ends while the pointed ends depolymerize very slowly. Once the polarity of filaments was determined as evidenced by the exclusive and rapid elongation from barbed ends, EGFP-MISP molecules were flowed in at a final concentration of 12.5 nM, and its binding position relative to the filament was monitored over time.

#### MISP-driven anchoring of F-actin

Biotin-anti-6xHis antibodies (SIGMA, MA1-21315-BTIN) were flowed in through a streptavidin-coated flow channel. Subsequently, EGFP-MISP molecules (harboring a 6xHis tag at the N-terminus) were flowed in at a final concentration of 12.5 nM, incubated for 5 minutes to allow for their immobilization. To monitor short filament capturing events, G-actin (500 nM) was mixed with TIRF polymerization buffer and polymerize for approximately 5 minutes before flowing in the reaction through the MISP-immobilized flow channel.

### ADP, ADP-Pi, and ATP-like filament preparation

F-Actin was generated as described earlier but was stabilized with unlabeled phalloidin (ThermoFisher, P3457) to allow for dilution below the critical concentration (< 100 nM). Phalloidin was also added at different time points of polymerization to generate F-actin in the ADP and ADP-Pi state as previously described.^34^ To note, phalloidin was included in all buffers throughout these assays to maintain the nucleotide state of filaments, especially in the case of the ADP-Pi condition.

#### ADP F-actin phalloidin stabilized

Rhodamine-biotin G-actin (1 µM) was polymerized at room temperature, and phalloidin (2 µM) was added at a 2:1 molar ratio. F-actin was further aged for 24 hours before conducting assays.

#### ADP-Pi F-actin phalloidin stabilized

Rhodamine-biotin G-actin (1 µM) was polymerized at room temperature in the presence of phalloidin (2 µM) at a 2:1 molar ratio, and used within the next 24 hours of preparation.

#### AMP-PNP F-actin phalloidin stabilized

Rhodamine-biotin G-actin (1 µM) was polymerized at room temperature in the presence of the nucleotide analog AMP-PNP (0.2 mM) and phalloidin (2 µM) at a 2:1 molar ratio, and used within the next 48 hours of preparation.

In all experiments, all different versions of actin filaments were flowed in into the TIRF channel, washed with TIRF buffer. Subsequently, EGFP-MISP in TIRF buffer was flowed in into the TIRF channel as imaging was in progress to determine dwell times.

### Actin bundling polarity assays

G-actin (500 nM) was mixed with EGFP-MISP (50nM) in TIRF polymerization buffer, and flowed in into an uncoated flow channel. To simplify quantification, two-filament bundling events were considered.

### Quantification and statistical analysis

All images were quantified and analyzed using the FIJI software package. Image registration from TIRF microscopy movies was performed using the StackReg plugin to correct for XY drifts when needed.

### MISP dwell time on phalloidin-stabilized F-actin

Line scans were drawn along movies of actin filaments longer than 2 µm and converted into kymographs. From these kymographs, the dwell time of multiple MISP binding events along the filament were measured and plotted in a histogram. Only finite MISP dwell times were considered for fitting purposes.

### MISP binding on polymerizing F-actin

The distribution of MISP binding events on filaments were calculated as follows: A single MISP binding event was considered positive if it remained stably bound to a single actin filament longer than 8 frames (i.e., 70 seconds). A line scan was drawn along the corresponding filament in the first frame of MISP binding to determine the filament length. The maximum MISP peak intensity was used as a reference point to determine its “distance from the pointed ends”.

### MISP apparent association rates

Rectangular ROIs (2 x 1.2 µm) were drawn along immobilized filaments (50 nM), which were used to measure EGFP-MISP (50 nM) fluorescence intensity over time. Intensity values were plotted and fitted with an exponential curve to estimate the apparent association rates.

### MISP-driven anchoring of single actin filaments

Anchoring events refer to events where a single polymerizing filament was immobilized by a single MISP puncta. A “pointed end anchoring event” was considered positive when the captured filament kept polymerizing from the opposite free end. A “barbed end anchoring event” was considered positive when the filament kept polymerizing from the captured end. Events that did not meet these criteria were considered as “side capturing events”.

### MISP-driven bundling of two actin filaments

We manually quantified the number of MISP-driven bundling events of two filaments by observing the direction of their growth before or after they overlapped. If filaments were already overlapping, their direction of growth was evident by the increase in the intensity of two filaments compared to one filament. To quantify the intensity of EGFP-MISP, we drew 0.5 µm x 2-pixel lines scans along 1- and 2-filaments at multiple time points. Values were then arbitrarily normalized to generate the plot.

### Statistical analysis

All statistical analysis was computed in PRISM v. 9.0 (GraphPad)

