## Supplementary figures and images for "Mitotic spindle positioning protein (MISP) is an actin bundler that senses ADP-actin and binds near the pointed ends of filaments"

### Morales and Tyska Supp Figs

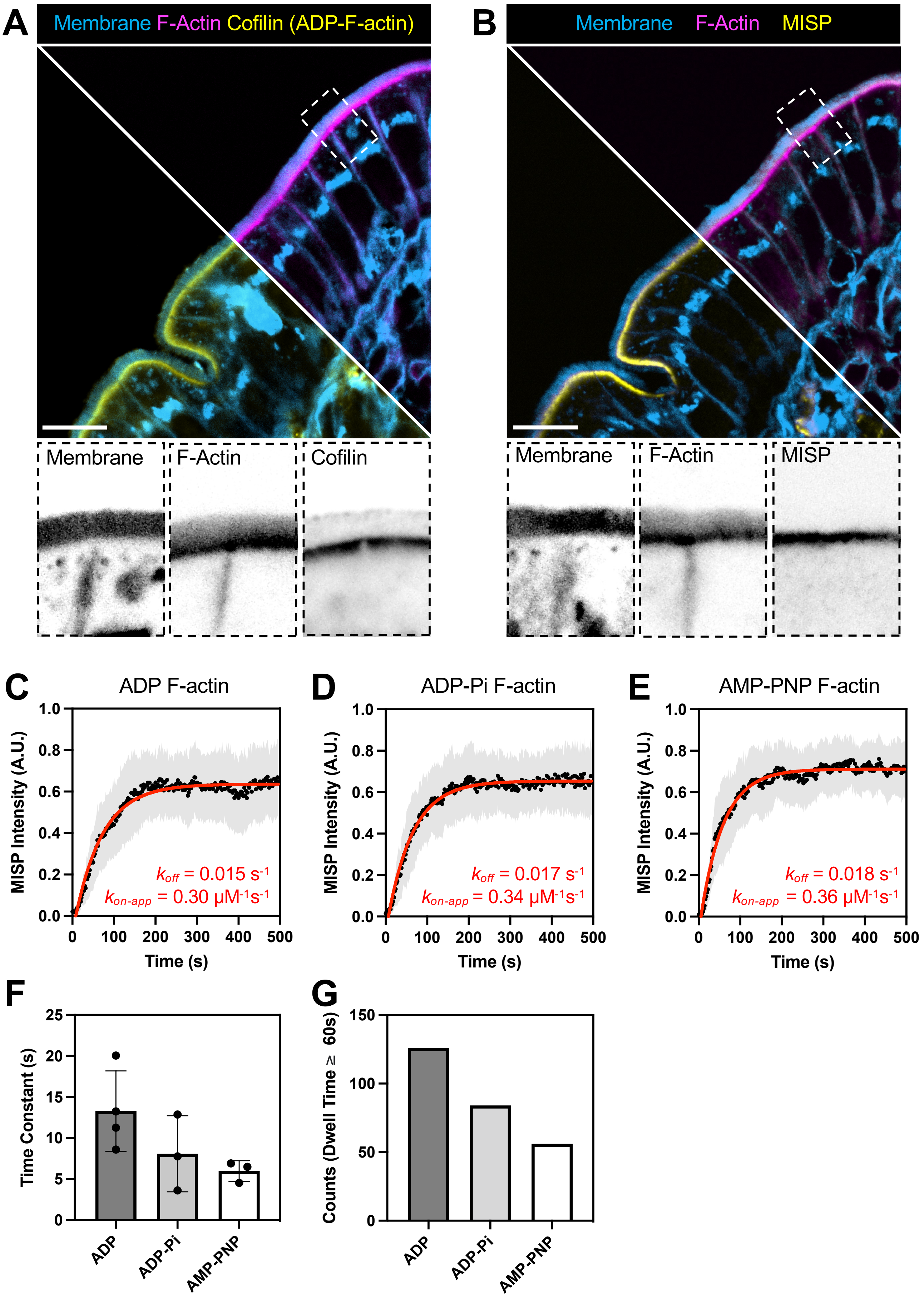

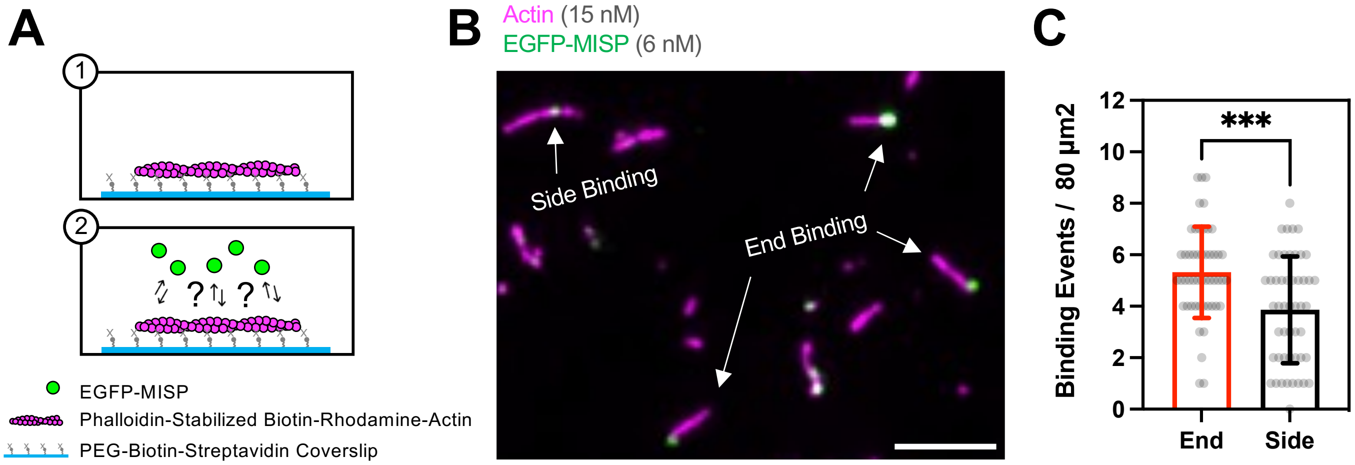

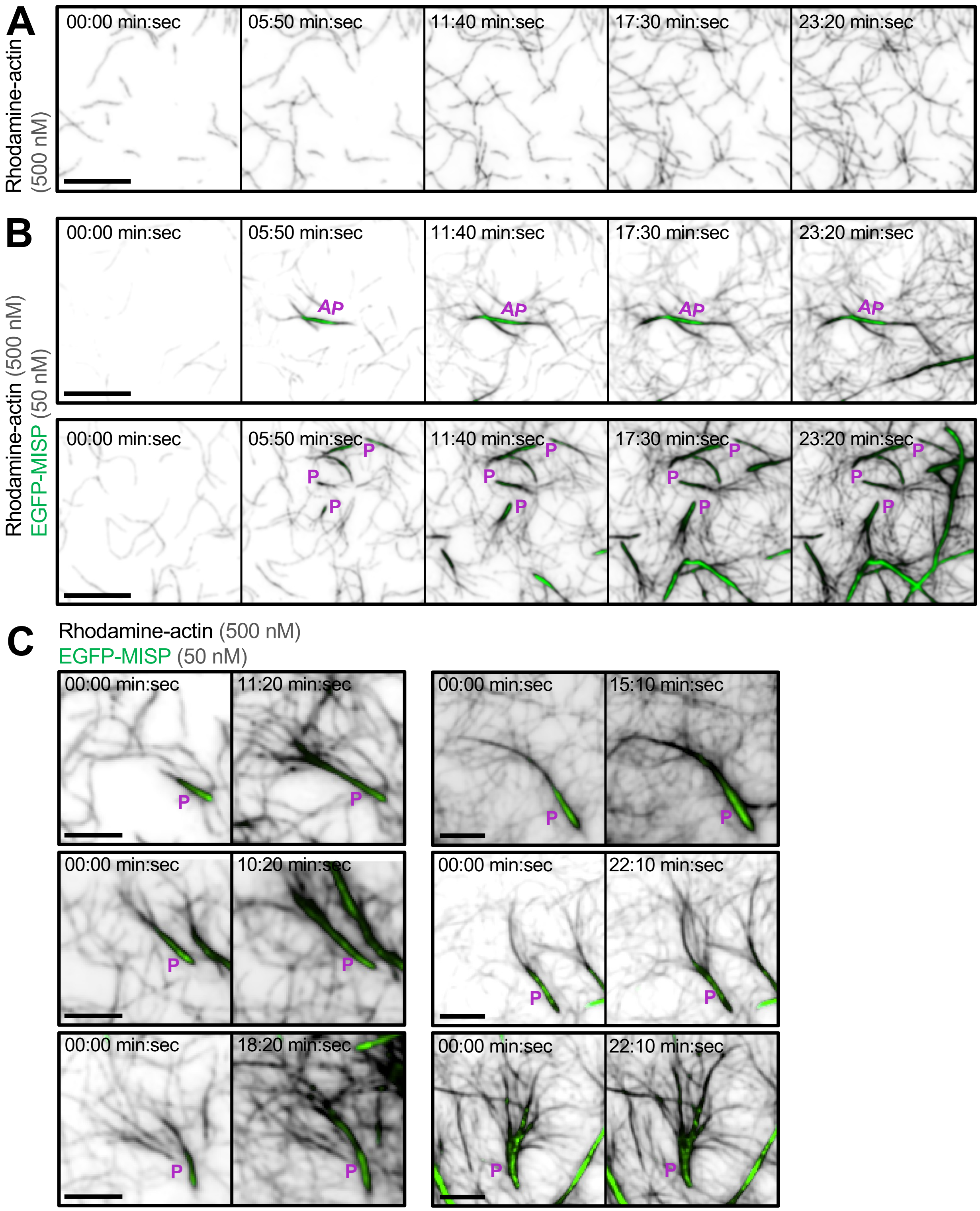
